# What drives the assembly of plant-associated protist microbiomes?

**DOI:** 10.1101/2020.02.16.951384

**Authors:** Kenneth Dumack, Kai Feng, Sebastian Flues, Melanie Sapp, Susanne Schreiter, Rita Grosch, Laura Rose, Ye Deng, Kornelia Smalla, Michael Bonkowski

## Abstract

In a field experiment we investigated the influence of the environmental filters soil type and plant species identity on rhizosphere community assembly of Cercozoa, a dominant group of (mostly bacterivorous) soil protists. The experiment was set up with two plant species, lettuce and potato, grown in an experimental plot system with three contrasting soils. Plant species (14%) and rhizosphere origin (vs. bulk soil) with 13%, together explained four times more variation in cercozoan beta diversity than the three soil types (7% explained variation in beta diversity). Our results clearly confirm the existence of plant species-specific protist communities. Network analyses of bacteria-Cercozoa rhizosphere communities identified scale-free small world topologies, indicating mechanisms of self-organization. While the assembly of rhizosphere bacterial communities is bottom-up controlled through the resource supply from root (secondary) metabolites, our results support the hypothesis that the net effect may depend on the strength of top-down control by protist grazers. Since grazing of protists has a strong impact on the composition and functioning of bacteria communities, protists expand the repertoire of plant genes by functional traits, and should be considered as ‘protist microbiomes’ in analogy to ‘bacterial microbiomes’.

**Highlight:** Microbiomes of rhizosphere protists are plant species-specific and tightly co-evolving with their bacterial prey, thereby extending and modifying the functional repertoire of the bacterial-plant symbiosis.

## Introduction

The assembly of specific subsets of the soil microbiota in the rhizosphere and root interior of plants has led to the characterization of plant species-specific ‘microbiomes’ (Berg *et al.*, 2014b; Hirsch and Mauchline, 2012; Lundberg *et al.*, 2012; Peiffer *et al.*, 2013). However, most attention has been given to microbial prokaryotes (Hacquard *et al.*, 2015; Martiny *et al.*, 2015; Müller *et al.*, 2016) and fungi (Philippot *et al.*, 2013; Porras-Alfaro and Bayman, 2011; Rodriguez *et al.*, 2009), while protists are virtually absent from models on plant microbiome assembly. A recent study of metatranscriptomics identified plant species-specific communities of bacterivorous Amoebozoa and Alveolata in the rhizospheres of pea, wheat and oat (Turner *et al.*, 2013). In addition, a metabarcoding study of Cercozoa (Rhizaria) found distinct subsets of bacterivorous protists associated with roots and leaves of *Arabidopsis thaliana* (Sapp *et al.*, 2017), demonstrating a close association of plants with specific protist communities.

These findings appear puzzling because protists, being major bacterivores in soil (Trap *et al.*, 2016), are thought to exert a large impact on the composition and functioning of rhizosphere bacterial communities (Bonkowski, 2004; Glücksman *et al.*, 2010; Jousset, 2012; Jousset *et al.*, 2008; Rosenberg *et al.*, 2009; Xiong *et al.*, 2018). The existence of plant species-specific protist communities indicates that each bacterial rhizosphere microbiome has an own adapted predator community, thus challenging current understanding of the regulation of rhizosphere processes.

Two major factors: soil type and plant species, determine the assembly of bacterial microbiota in the rhizosphere of plants (Berg and Smalla, 2009; Haichar *et al.*, 2008; Schreiter *et al.*, 2014a). Soil type with its specific physical and chemical properties determines the resident microbial community (Girvan *et al.*, 2003; Sessitsch *et al.*, 2001; Ulrich and Becker, 2006), from which plant species recruit specific subsets of rhizosphere microbiota due to the growth-limiting carbohydrates and distinct metabolite profiles provided in root exudates (Baetz and Martinoia, 2014; Jones *et al.*, 2009; Sasse *et al.*, 2018; van Dam and Bouwmeester, 2016).

Consistent with these studies, the greenhouse experiment by Sapp *et al.* (2017) demonstrated a strong structuring effect of soil type on the cercozoan protist communities of *Arabidopsis thaliana*. However, a rigorous testing of the existence of plant species-specific associations of protist microbiota, and their modification by soil conditions can only be achieved in field experiments. In order to verify the existence of plant-specific protist ‘microbiomes’ under natural conditions, we applied the cercozoan primers used by Sapp *et al.* (2017) in a factorial field experiment with two plant species, lettuce and potato. These plants were grown in close proximity to one another in an experimental field plot system with three contrasting soils (see Schreiter *et al.*, 2018) to obtain a robust measure of the factors influencing protist rhizosphere microbiomes. Variance partitioning allowed the quantification of the influence of the environmental filters (soil type and plant species) on community assembly of Cercozoa. We further performed network analyses of Cercozoa and their co-occurrence with their potential bacterial prey on lettuce and potato (Schreiter *et al.*, 2018) to better characterize bacteria-protist relationships.

## Material and Methods

### Field experiments and sampling

A field experiment was set up with lettuce (*Lactuca sativa* L.; cv. Tizian, Syngenta, Bad Salzuflen, Germany) and potato (*Solanum tuberosum* L.; cv. Arkula, Norika GmbH, Groß Lüsewitz, Germany). The plants were grown in three different soil types in a unique experimental plot system in independent experimental units at the Leibniz Institute of Vegetable and Ornamental Crops (IGZ, Großbeeren, Germany, 52° 33’ N, 13° 22’ E). Two units were used in this study, each containing three soil types characterized as Arenic-Luvisol (diluvial sand, DS), Gleyic-Fluvisol (alluvial loam, AL), and Luvic-Phaeozem (loess loam, LL) sharing the same climatic conditions and each unit the same crop history for more than 10 years (Schreiter *et al.*, 2014a). The soil types were arranged in separate blocks (one per soil type) with 24 plots of 2 m × 2 m in size and a depth of 75 cm. Potato and lettuce were planted in a randomized design in experimental plots of separate experimental units on 15^th^ June and 3^rd^ July 2012, respectively. Seed potato tubers were planted 30 cm apart within a row and with an intra-row distance of 65 cm (21 tubers per plot), while lettuce was planted with a within-row and intra-row distance of 30 cm between plants (36 plants per plot). Each plant treatment and soil type treatment was replicated four times. Rhizosphere soil samples of lettuce were collected two weeks after planting, rhizosphere soil samples of potato were taken seven weeks after planting. More details on the experimental design can be found in (Schreiter *et al.*, 2018).

### Sample processing

The roots of two potato plants or three lettuce plants per plot were pooled to reduce intra-plot variability. Adhering soil was removed by a short root washing step and afterwards the roots were cut into pieces of 1 cm, mixed and 5 g of roots were treated three times by a Stomacher 400 Circulator (Seward Ltd., Worthing, United Kingdom) as described in Schreiter *et al.* (2018). The portion of soil still sticking to the root was denoted as rhizosphere soil. Bulk soil samples were taken between planted rows (Schreiter *et al.*, 2014b). Sample processing, DNA extraction followed (Schreiter *et al.*, 2014a).

### Molecular analyses

Sequencing of the bacterial 16S SSU rDNA V3/V4 region, including PCR reactions and pre-filtering was conducted by the Biotechnology Innovation Center (BIOCANT, Cantanhede, Portugal) on a 454 Roche sequencing platform as described in Schreiter *et al.* (2014b). Subsequently sequences of less than 200 bp length were excluded and clustered at 97% in mothur v.3.9 (Schloss *et al.*, 2009) to create operational taxonomic units (OTUs). OTUs were verified with UCHIME (Edgar *et al.*, 2011) as implemented in mothur and identified using BLAST+ (Camacho *et al.*, 2009) with the SILVA database as a reference (Pruesse *et al.*, 2007).

PCRs of the cercozoan community were conducted in two steps. In the first PCR, the forward primers S616F_Cerco and S616F_Eocer were mixed in the proportions of 80% and 20%, and used with the reverse primer S963R_Cerco (Fiore-Donno *et al.*, 2018). One µl of ten times diluted DNA were used as a template for the first PCR and 1 µl of the resulting amplicons were used as a template for a following semi-nested PCR. We employed the following final concentrations: Dream Taq polymerase (Thermo Fisher Scientific, Dreieich, Germany) 0.01 units, Thermo Scientific Dream Taq Green Buffer, dNTPs 0.2 mM and primers 1 µM. The conditions were set to an initial denaturation step at 95°C for 2 min, 24 cycles at 95°C for 30 s, 50°C for 30 s, 72°C for 30 s; and a final elongation step at 72°C for 5 min. The second PCR was conducted with barcoded primers (see Fiore-Donno *et al.*, 2018). All PCRs were conducted twice to reduce the possible artificial dominance of few amplicons by PCR competition, and then pooled.

A mock community with known species richness of diverse cultivated cercozoan taxa was run in parallel to assist the fine-tuning of the bioinformatics pipeline as described in Fiore-Donno et al. (2017). The amplicons were checked by electrophoresis and 25 µl of each pooled PCR product were purified and normalized using SequalPrep Normalization Plate Kit (Invitrogen GmbH, Karlsruhe, Germany). We then pooled the samples and the mock community and proceeded for a single library preparation. Library preparation and paired-end MiSeq sequencing with the MiSeqv3 2×300 bp kit were carried out by the Cologne Center for Genomics (CCG).

Paired reads were assembled using mothur v.3.9 (Schloss *et al.*, 2009) allowing one difference in the primers, no difference in the barcodes, no ambiguities, no mismatches greater than three and removing assembled sequences with an overlap <200 bp. Reads were sorted into samples according to the barcodes (Table S1). The quality check and removal/cutting of low-quality reads were conducted with the default parameters. Using BLAST+ (Camacho *et al.*, 2009) with an e-value of 1e^−50^ and keeping only the best hit, sequences were identified in the PR2 database (Guillou *et al.*, 2013) and non-cercozoan sequences were removed. Chimeras were identified using UCHIME (Edgar *et al.*, 2011) as implemented in mothur with a penalty for opening gaps of −5 and a template for aligning operational taxonomic units (OTUs, V4 region of 78 cercozoan taxa, see Fiore-Donno *et al.*, 2018). Sequences were clustered using VSEARCH v.1 (Rognes *et al.*, 2016), with abundance-based greedy clustering (agc) and a similarity threshold of 97% as indicated by analyzing the mock community. A cutoff was determined by the mock community and OTUs representing less than 4% of reads were deleted. Pyrosequence data were deposited at the European Nucleotide Archive under the study accession number ERS4306420.

### Statistical analyses

All statistical analyses and data visualizations, except networks, were conducted in R version 3.1.1 (R Core Team, 2014). First a table of the frequency of OTUs for each sample was generated and normalized by dividing by the total number of OTUs. We calculated Shannon diversity and Pielou’s evenness to compare cercozoan bulk soil and rhizosphere soil communities. We further used non-metric multidimensional scaling (NMDS) based on Bray-Curtis dissimilarities to visualize the community structure between treatments using the normalized OTU abundance matrix generated as described above. Permutational Multivariate Analysis of Variance (PERMANOVA) (Anderson, 2001) using Bray-Curtis dissimilarity was employed to test differences in Cercozoa community assembly across treatments.

We used variance partitioning analysis (varpart function in the vegan package, version 2.3-5 in R) to quantify the variance in beta-diversity of Cercozoa explained by soil types, rhizosphere vs. bulk soil and plant species (lettuce vs. potato) and their combined effects. The function uses adjusted R^2^ to assess the partitions explained by the explanatory variables and their combinations (Peres-Neto *et al.*, 2006). We ran permutation tests to test the significance of all constraints simultaneously (Oksanen *et al.*, 2015). All tests and plots were performed using the vegan package (Oksanen *et al.*, 2015) and each test was permuted 999 times.

#### Network analyses

Network analyses were performed to investigate if co-occurrence deviated from random patterns, and to assess the complexity of potential interactions between bacteria and cercozoan protists in the lettuce and potato rhizospheres. Co-occurrence analyses were performed using the molecular ecological network analysis pipeline (MENAP, http://ieg4.rccc.ou.edu/mena/) (Deng *et al.*, 2012; Zhou *et al.*, 2011). For each network, only the OTUs present in more than nine samples were kept to calculate a Spearman rank correlation matrix without log-transformation, and then the entire network was generated with a finest threshold (0.85) according to random-matrix theory (RMT) judgements.

The network topological features were calculated in MENAP. Eleven features were evaluated: 1) total number of nodes (N); 2) numbers of total links; 3) intra-domain links (between only bacterial or only cercozoan nodes); 4) inter-domain links (between bacterial and cercozoan taxa); 5) connectance, i.e. the number of established links relative to the number of expected links; 6) betweenness centrality characterizing the number of pathways that go through a particular OTU when it is between a pair of other OTUs; 7) whether connections (k) per node (N) followed a power law (N(k)~k^−^□).(R^2^ of power law); 8) the average degree measuring the average connectivity of OTUs in a network; 9) the average path distance as a measure of network diameter measuring the average of the distances between each pair of nodes in the network; 10) the average clustering coefficient describing the grouping of closely connected subsets of nodes into highly connected groups or ‘cliques’; and 11) modularity identifying separate modules of connected nodes at the network scale (Delmas *et al.*, 2019).

Based on modularity results, the topological roles of nodes could be assigned into four different ecological categories by within-module connectivity (*z*) and among-module connectivity (*P*) (Guimera and Amaral, 2005): peripheral nodes (*z* ≤ 2.5, *P* ≤ 0.62) that have few links to other nodes both within and among modules, module hubs (*z* > 2.5, *P* ≤ 0.62) that were highly connected to nodes within modules, connectors (*z* ≤ 2.5, *P* > 0.62) that were highly connected to nodes among modules, and network hubs (*z >* 2.5, *P >* 0.62) that act as both connectors and module hubs (Olesen *et al.*, 2006). Furthermore, we used additional indicators to assign keystone taxa and made comparisons between lettuce and potato, i.e. nodes with maximum betweenness (characterizing the number of pathways that go through a particular OTU when it is between a pair of other OTUs) and nodes with maximum node degree (number of links).

In order to focus on the inter-domain associations, the links between Cercozoa and bacteria and the nodes affiliated to these links were extracted to generate sub-networks. For visualization the nodes were grouped at the family level and the sum of either positive or negative correlations was displayed as width of inter-family edges in the network graph. Intra-family links were ignored. The network graphs were visualized with Cytoscape 3.3.0 software (Shannon *et al.*, 2003).

## Results

We identified 249 cercozoan OTUs out of an initial 7,335,204 sequences that passed our quality filters (see Methods). Rarefaction curves show that sequencing depth was sufficient to reach saturation (Fig. S1). A database with the abundance of each OTU per site and its taxonomic assignment is provided as supplement (Table S2). The dominant cercozoan groups were Glissomonadida and Cercomonadida, representing mostly small flagellates and amoeboflagellates (Fig. 1).

**Fig 1.**
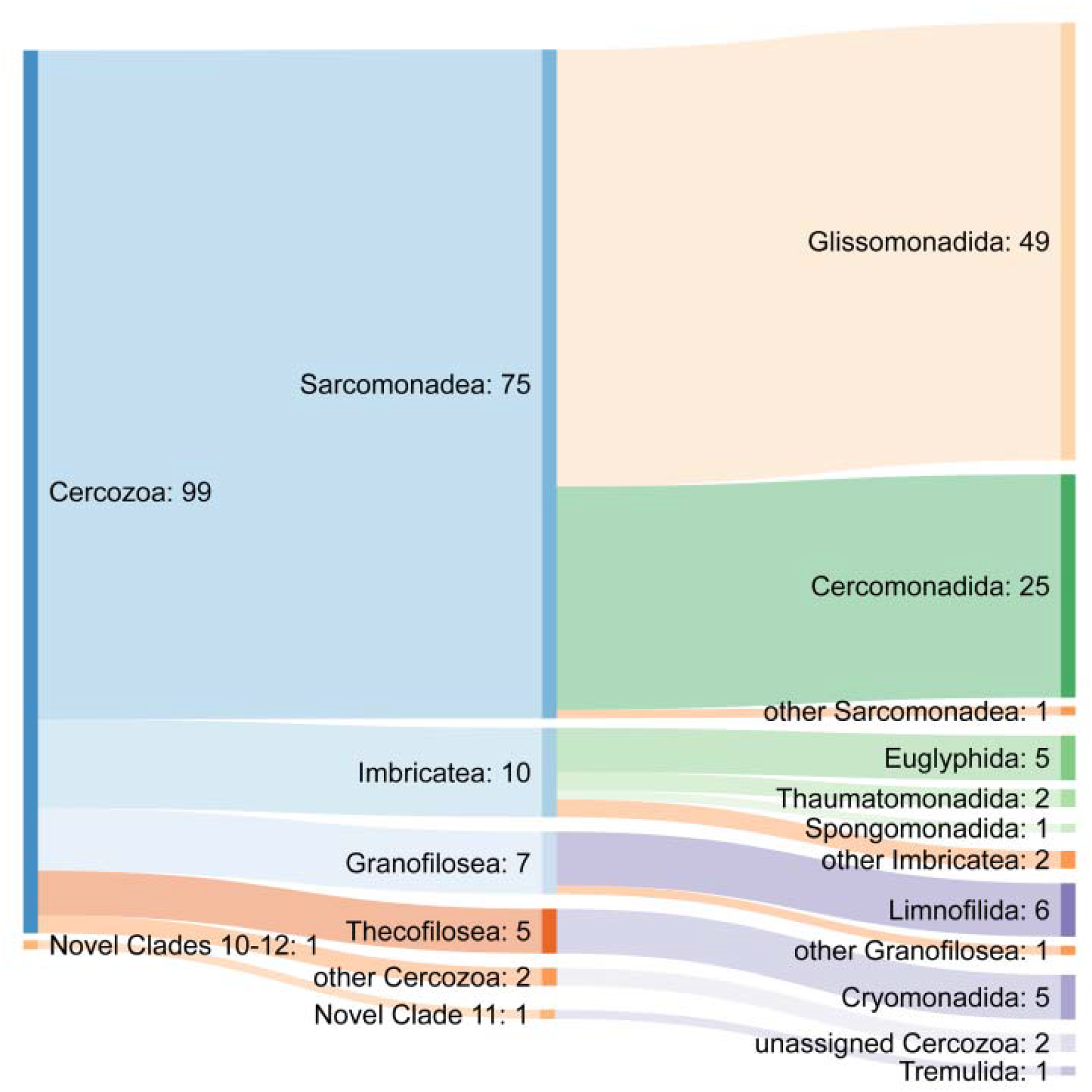
Sankey diagram showing the relative contribution of OTUs to the taxonomic diversity. Taxonomical assignment was based on the best hit by BLAST. From left to right, names refer to phyla (Cercozoa, Endomyxa), class (ending -ea), and orders (ending -ida). “Others” refer to sequences that could either not be assigned to the next lower-ranking taxon or made up less than 1% of cercozoan diversity. Numbers are percentages of sequence abundance.

Cercozoan alpha diversity was significantly lower in rhizosphere soil (H_lettuce_ 3.78; H_potato_ 3.76) than in bulk soil (H_bulk_ 4.3; F_[2,117]_ = 52.53; *P*<0.001). The same was true for cercozoan evenness in rhizosphere (J_lettuce_ 0.70; J_potato_ 0.70) compared to bulk soil (J_bulk_ 0.78; F_[2,117]_ = 42.37; *P*<0.001, Fig. S2). All Cercozoa considered in the analyses were bacterivores. Plant parasitic Endomyxa contributed less than 1% to cercozoan OTUs and were not included.

### Soil type and plant species dependent assembly of cercozoan rhizosphere communities

The composition of bulk soil communities of Cercozoa were different in loam compared to sand and loess (PERMANOVA R^2^ = 0.33, F_2,21_ = 5.05, *P* = 0.001, Fig. 2). However, the plant rhizosphere exerted a particularly strong effect on the community structure of Cercozoa. Cercozoan community assembly in the rhizosphere was influenced by soil type (PERMANOVA R^2^ = 0.15, F_2,89_ = 15.73, *P* = 0.001) and strongly dependent on plant species identity (PERMANOVA R^2^ = 0.31, F_1,89_ = 64.99, *P* = 0.001). A significant interaction of soil type and plant species (PERMANOVA R^2^ = 0.11, F_2,89_ = 11.25, *P* = 0.001) reflects the fact that soil type had a stronger effect on the assembly of cercozoan communities under lettuce than under potato (Fig. 2).

**Fig. 2.**
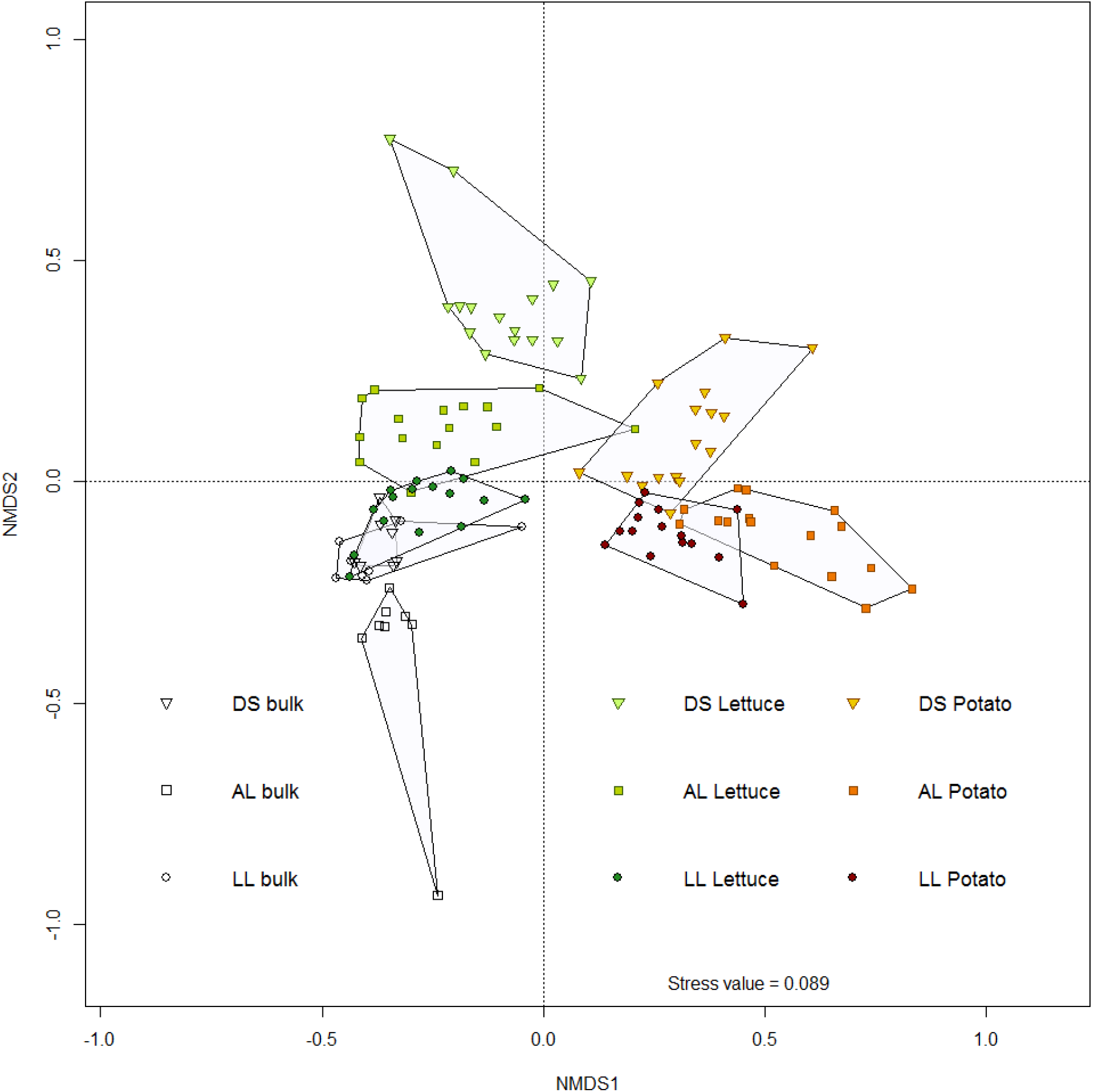
Non-metric multidimensional scaling (NMDS) of Bray-Curtis dissimilarities among cercozoan communities for lettuce (green colors) and potato (brown colors) separated by soil type with diluvial sand (DS, triangle), alluvial loam (AL, square), and loess loam (LL, circle) (PERMANOVA R^2^ = 0.62, F_8,110_ = 22.14, *P* = 0.001). NMDS stress value was 0.089.

Variance partitioning (Fig. 3) allowed the quantification of explained variation in cercozoan beta diversity by the three soil types (6.7%; F = 9.10, *P* = 0.001), by plant species identity (14.4%; F = 20.15, *P* = 0.001) and by differences between rhizosphere and bulk soil (13.0%; F = 17.96, *P* = 0.001). Thus rhizosphere origin and differences between both plant species together explained four times more variation of cercozoan community composition than differences in beta diversity between the three contrasting soil types.

**Fig. 3.**
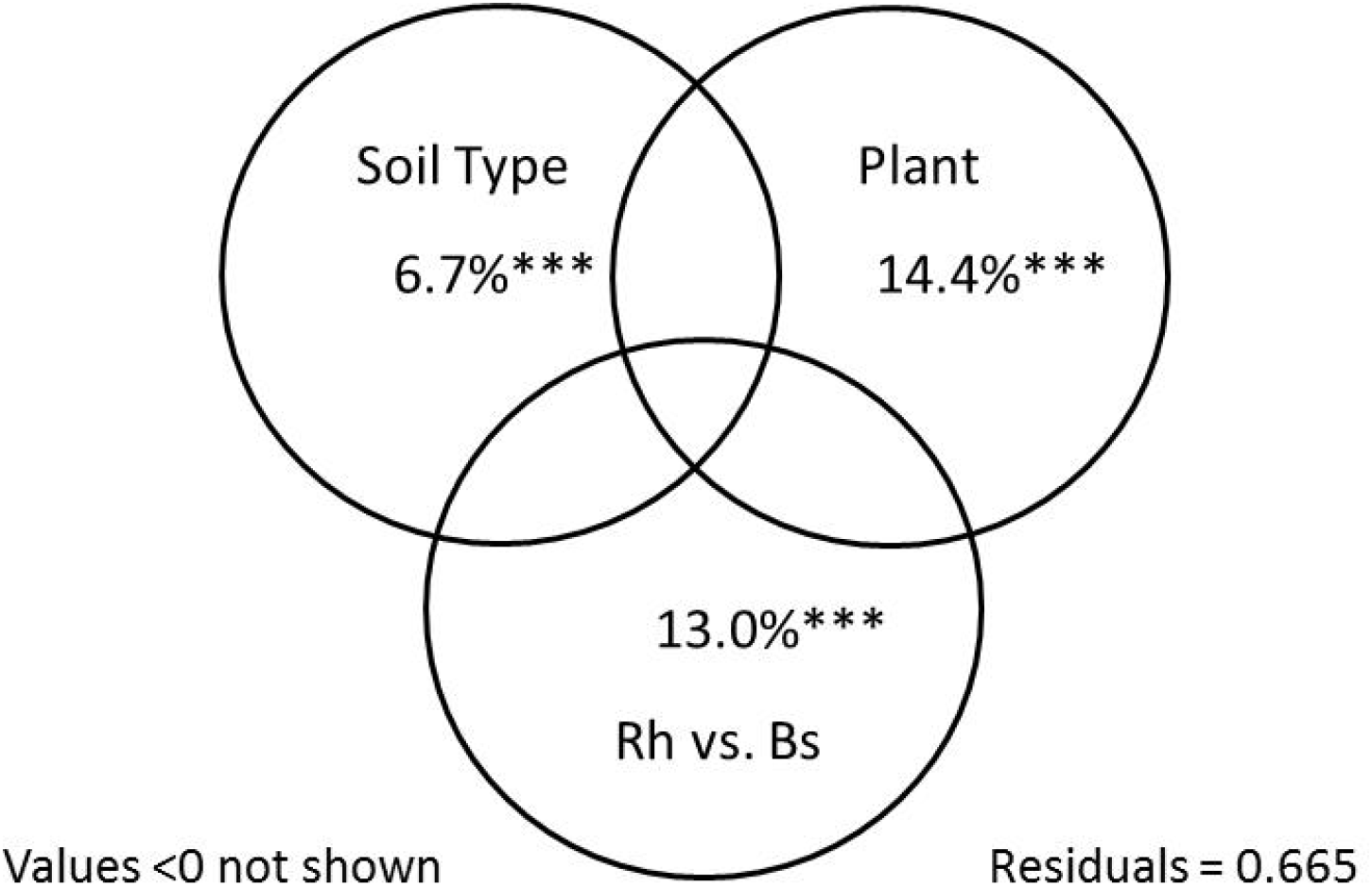
Partitioning of variance explained in beta diversity of cercozoan communities by three different soil types (Soil Type), plant species identity (Plant) and rhizosphere versus bulk soil (Rh vs. Bs). Residuals of unexplained variance were 0.665, *** indicates P < 0.001.

### Modular networks with scale-free, small world architecture

To better understand potential bacteria-protist interactions, we performed co-occurrence analyses of the 249 cercozoan OTUs and 8203 bacterial OTUs in the rhizospheres of lettuce and potato. The networks for lettuce and potato showed non-random topologies with a modular structure (Fig. 4, Table 1). The network connectivity was uneven and followed a power law, characteristic of scale-free networks with a topology of many nodes with few connections and some highly connected nodes (i.e. hub taxa), having densely positioned nodes within modules (small values of average path distance, Table 1). The networks for lettuce and potato contained 44 and 45 modules, respectively. Compared to random networks, the modularity of the lettuce and potato networks (MOD/MOD_*random*_) was elevated (1.28 for lettuce and 1.34-fold for potato; Table 1, one sample student’s *t* test, *P*<0.001). Clustering of the lettuce networks was 10 times higher than that of random networks and 15 times higher for potato compared to random networks (avgCC/avgCC_*random*_), indicating clustered, and highly correlated sub-networks. Average path distance (APD/APD_*random*_, a measure of distance between nodes indicative of network size), was elevated by a factor of 1.43 in the lettuce network and 1.6 in the potato network compared to random networks (Table 1, one sample student’s *t* test, *P*<0.001). This means that the networks could be sub-divided into modules with clustered, highly interconnected nodes characteristic of a small world architecture. The highly connected nodes (i.e. maximum degree) could be identified as a cercozoan *Neoheteromita globosa* in lettuce and a bacterial *Sphingomonas* in potato networks (Table S4). All three measures (modularity, avgCC, and APD) were slightly, but significantly higher for potato than lettuce networks (student’s *t* test, *P*<0.001). Taken together, both bacteria-cercozoa networks exhibited a scale-free, small world architecture (Table 1).

**Table 1:** Topological features of empirical lettuce and potato bacteria-Cercozoa rhizosphere networks and of associated random networks generated by randomly rewiring all nodes and links 100 times. The following features are reported: similarity threshold; total number of nodes (total nodes); number of nodes consisting of protists (Protists); number of nodes consisting of bacteria (Bacteria); number of total links (total links); number of links only between bacterial taxa or cercozoan taxa (Intra-domain links); number of links between bacterial taxa and cercozoan taxa (Inter-domain links); connectance; and betweenness centrality; the proportion of variance explained under the assumption that the number of connections per node followed a power law function (R^2^ of power law); average number of connections (average degree, avgK); average clustering coefficient (avgCC); average distance between nodes (average path distance, APD); and number of network modules (Modularity, MOD).

|  | Features | Lettuce | Potato |
| --- | --- | --- | --- |
| Empirical networks | Similarity threshold | 0.85 | 0.85 |
|  | Total nodes | 276 | 345 |
|  | Protists | 130 | 164 |
|  | Bacteria | 146 | 181 |
|  | Total links | 441 | 556 |
|  | Intra-domain links | 111 | 338 |
|  | Inter-domain links | 310 | 218 |
|  | Connectance (Con) | 0.536 | 0.539 |
|  | Betweenness Centrality (BC) | 0.136 | 0.133 |
| | $R^2$ of power law<sup>b</sup> | 0.989 | 0.975 |
|  | Average degree (avgK) | 3.196 | 3.223 |
|  | Average clustering coefficient (avgCC)<sup>c</sup> | 0.181<sup>d</sup> | 0.210<sup>d</sup> |
|  | Average path distance (APD)<sup>c</sup> | 6.117<sup>d</sup> | 7.152<sup>d</sup> |
|  | Modularity (MOD)<sup>c</sup> | 0.732 (44)<sup>d</sup> | 0.779 (45)<sup>d</sup> |
| Random networks<sup>a</sup> | Average clustering coefficient (avgCC) | 0.018 ± 0.006 | 0.014 ± 0.005 |
|  | Average path distance (APD) | 4.284 ± 0.071 | 4.462 ± 0.065 |
|  | Modularity (MOD) | 0.573 ± 0.008 | 0.580 ± 0.007 |

- a. Random networks were generated by randomly rewiring all nodes and links 100 times b. Test if the number of connections (k) per node (N) followed a power law ( $N(k) \sim k^{-1}$ ). c. Significant difference ( $P < 0.001$ ) in avgCC, APD and MOD of empirical networks compared to random networks for both lettuce and potato, based on one sample student's *t* test. d. Significant difference ( $P < 0.001$ ) in avgCC, APD and MOD between lettuce and potato using Student's *t* test.

**Fig. 4.**
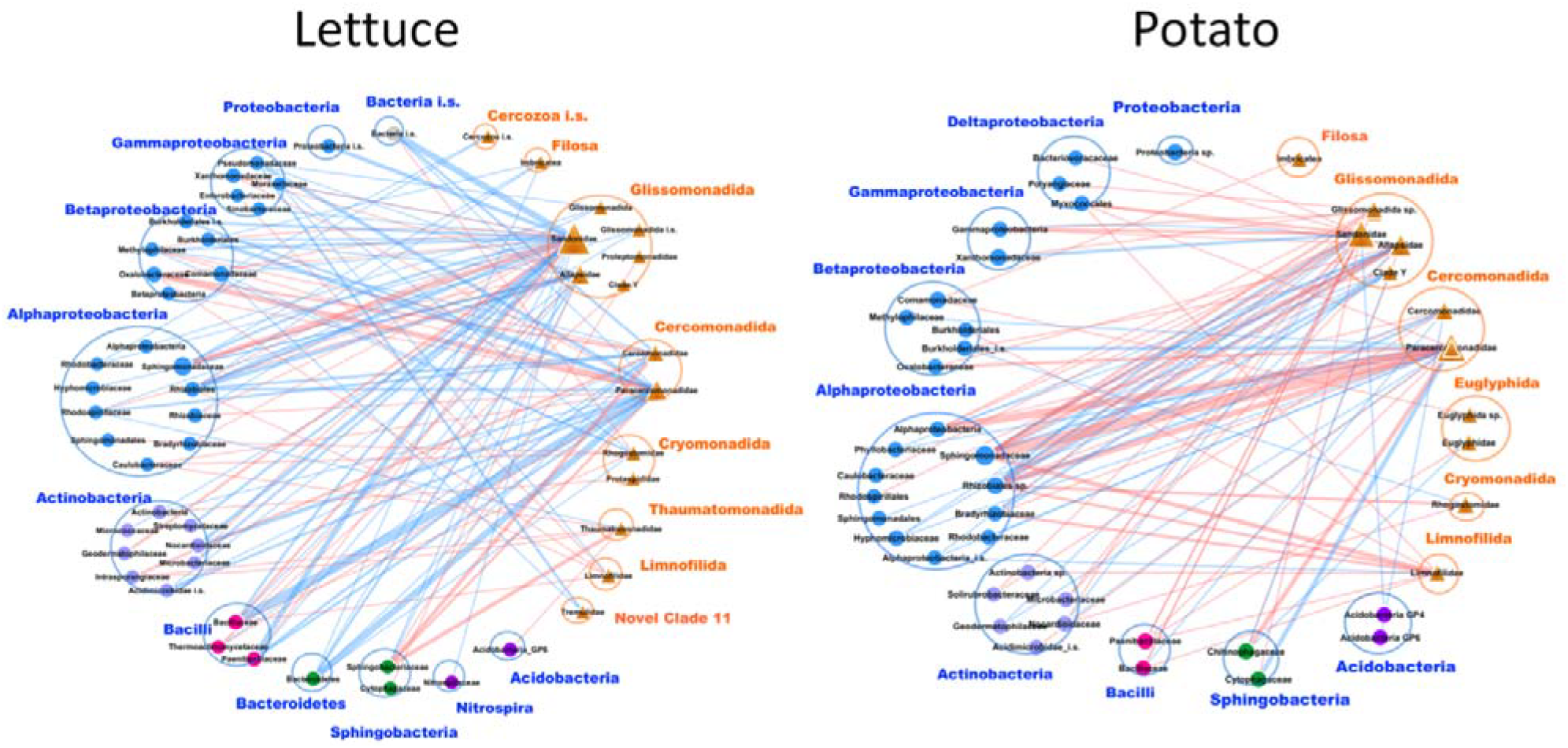
Microbial co-occurrence networks based on correlation analysis of bacteria (circles) and cercozoan protists (orange triangles) for lettuce (left) and potato (right). The relative abundance of bacteria and Cercozoa is represented on family level by the size of nodes. Node colors were mapped to the phylum level. A connection shows the union of negative or positive co-occurrence between bacterial communities and cercozoan communities on OTU level. Positive and negative co-occurrences are indicated by blue and red edges, respectively, whereas the edge widths indicate the proportion of correlations among OTUs between two families of bacteria and protists. Nodes were clustered on family level based on their current taxonomy and loops that indicate co-occurrence relationships of microbial species of the same trophic level or family were removed.

The most striking difference between bacteria-Cercozoa networks for the two plant species were the presence of mainly positive co-occurrences for lettuce and mainly negative co-occurrences for potato (Fig. 4). Paracercomonadidae formed a highly connected node in both networks, however it showed positive co-occurrences with Bacillaceae and Bacteroidetes for lettuce, while its associations with bacteria for potato were mostly negative, in particular with Sphingomonadaceae, Rhizobiales and Alphaproteobacteria. Nodes with maximum betweenness were occupied by protists belonging to uncharacterized Limnofilidae in lettuce and Allapsidae in potato (Table S4).

The module hubs for potato were mainly Cercozoa (an unclassified Allapsidae belonging to the as yet undescribed Group Te, and undescribed members of Imbricatea and Clade Y in Glissomonadida) and the connectors were bacteria (Rhizobiales) (Fig. S3). For lettuce, the module hubs mainly belonged to bacteria and the connectors included both bacteria belonging to Burkholderiales and Cercozoa (*Paracercomonas compacta*, and an unclassified Limnofilidae, Fig. S3).

## Discussion

Bacterivorous flagellates, amoeboflagellates and testate amoebae dominated the cercozoan community (Fig. 1). Exemplified by Cercozoa, our results clearly confirm the existence of plant species-specific ‘protist microbiomes’ of bacterivores in the rhizosphere of field grown plants as postulated by Sapp *et al.* (2017). Bulk soil contained a higher diversity and evenness of cercozoan OTUs compared to the rhizospheres of lettuce and potato, which corresponds to findings on bacterial microbiota (Shi *et al.*, 2015). The reduced protist diversity and the four-fold stronger combined effect on cercozoan community composition by the rhizosphere (i.e. its specific modification relative to bulk soil) and plant species identity compared to soil type (Fig. 3) reveal that the plant rhizosphere is a strong habitat filter for protist community assembly.

The bacterial taxa in our networks have been identified as typical members of lettuce and potato ‘core microbiomes’ (Cardinale *et al.*, 2015; Mitter *et al.*, 2016; Schreiter *et al.*, 2014a; Schreiter *et al.*, 2018). The term ‘microbiome’ *sensu stricto* denotes the microbial genes encoding specific traits supplementing the plant genome by microbial functions such as nutrient provision or pathogen defense (Berg *et al.*, 2014a; Sánchez-Cañizares *et al.*, 2017; Vandenkoornhuyse *et al.*, 2015). Correspondingly, the ‘protist microbiome’ supplements the plant genome by beneficial protist functions. These may include the provision of growth-limiting nutrients to plants (Bonkowski and Clarholm, 2012; Bonkowski *et al.*, 2000; Ekelund *et al.*, 2009) and associated mycorrhiza (Bonkowski *et al.*, 2001; Bukovská *et al.*, 2018; Jentschke *et al.*, 1995; Koller *et al.*, 2013a; Koller *et al.*, 2013b), the direct control of plant pathogenic fungi (Chakraborty *et al.*, 1983), or enhancing the expression of bacterial biocontrol genes and metabolites against plant pathogens (Jousset and Bonkowski, 2010; Jousset *et al.*, 2009; Jousset *et al.*, 2010).

Lettuce and potato specific cercozoan ‘microbiomes’ of bacterivorous protists however appear to contradict these earlier studies showing that protist communities were shaped by plants or their associated communities of rhizobacteria, instead of rhizosphere bacterial communities being shaped by the grazing pressure of protists (Jousset *et al.*, 2010; Jousset *et al.*, 2008; Rosenberg *et al.*, 2009; Saleem *et al.*, 2012).

This raises the question on the mechanisms underlying the plant species-specific assembly of protists. For rhizosphere bacteria a bottom-up regulation through resource supply from roots is seen as the major driver of community selection (Bakker *et al.*, 2015; Sasse *et al.*, 2018). The composition of root exudates, by providing a crucial energy source for soil microorganisms (Kuzyakov and Blagodatskaya, 2015) and containing secondary metabolites as microbial attractants or chemical deterrents have been suggested to select for the plant species specific microbiomes (Guyonnet *et al.*, 2018; Sasse *et al.*, 2018).

Analogously, secondary metabolites of bacteria may shape the assembly of protist predators in the rhizosphere of plants. Bacterivorous protists trigger immediate changes in bacterial chemical defense (Flues *et al.*, 2017; Jousset and Bonkowski, 2010; Jousset *et al.*, 2006; Jousset *et al.*, 2010). Defense is energetically costly, causing inequalities due to competitive trade-offs in the growth-defense balance of bacterial communities (Jousset *et al.*, 2009).

Accordingly, shifts in predation pressure sorts out winners and losers among bacteria, resulting in a functional and taxonomic remodeling of bacterial communities (Flues *et al.*, 2017; Glücksman *et al.*, 2010; Rosenberg *et al.*, 2009; Xiong *et al.*, 2018). Overall, grazing-resistant bacterial taxa which exhibit targeted allelopathy against eukaryotes are favored in soil systems (Arp *et al.*, 2018; Jousset, 2012; Jousset *et al.*, 2008; Matz and Kjelleberg, 2005; Mazzola *et al.*, 2009). This again may have important consequences for plant performance, not only because some of these metabolites directly or indirectly influence root growth (Brazelton *et al.*, 2008; Combes-Meynet *et al.*, 2011), but because the same defense compounds ward off microbial competitors, including fungal and bacterial plant pathogens (Arp *et al.*, 2018; Meyer *et al.*, 2009; Ramette *et al.*, 2011; Russell *et al.*, 2014). Accordingly, the resulting communities of rhizosphere bacteria have been shown to express enhanced biocontrol activity, indicating increased reliability of microbiome function (Jousset *et al.*, 2011; Rosenberg *et al.*, 2009; Weidner *et al.*, 2016).

In correspondence with this hypothesis, our network analyses indicate non-random co-occurrences of Cercozoa and bacteria at the family level (see superscript^**c**^, Table 1). Scale-free networks exhibit specific mechanisms of self-organization, where highly connected nodes acquire links at a higher rate than those that are less connected. This leads to the emergence of a few highly connected hubs (Barabási, 2009; Montoya *et al.*, 2006; Watts and Strogatz, 1998).

In a food web context, the constant release of root exudates favoring specific rhizosphere bacteria and reciprocal specialized predators could result in the accumulation of co-evolved subsets of rhizosphere microbiota, leading to positive co-occurrences as seen for lettuce. Over the longer term, the accumulation of allelopathic metabolites may restrict the activity of protists (Foissner, 1987; Jousset, 2012; Jousset *et al.*, 2006), and could lead to negative co-occurrences similar to those seen for potato (Fig. 4).

‘Small world’ topologies characterize highly interconnected sub-networks which are resilient to perturbations, because random losses of node species may be easily compensated by links to other nodes, except if a key node is affected (Albert *et al.*, 2000; Montoya *et al.*, 2006). In this study, such ‘keystone taxa’ were identified as the cercozoan amoeboflagellate *Neoheteromita globosa* in lettuce and a *Sphingomonas* bacterium in potato. A pronounced edge width between bacteria and *Paracercomonas* and Sandonidae in both networks suggests a strong impact of these protist taxa on microbiome structure. The pronounced edge width may further indicate a certain degree of functional redundancy among *Paracercomonas* and Sandonidae, which could act as ‘trophic species’ where phylogenetically related predators may exhibit similar prey preferences. If true, such functional redundancy may contribute to the stability and self-organization of food-web relationships in the rhizosphere.

However, trade-offs may arise because the performance of cercozoan species differs in response to the composition of bacterial assemblages (Flues *et al.*, 2017; Glücksman *et al.*, 2010; Xiong *et al.*, 2018). In a key experiment, manipulating the diversity of protist predators and their bacterial prey, Saleem *et al.* (2013) identified the synergistic exploitation of bacterial prey by predator complementarity as main driver of protist community performance. Thus prey-predator matching may lead to an optimization and functional stabilization of these interactions.

Overall, this study laid the foundation of a number of testable new hypotheses on microbiome assembly and functioning. Most importantly, our results suggest ripple effects of root metabolites via bacteria to the next trophic level. A dynamic feedback of rhizosphere bacteria communities on protist community assembly and vice versa has far reaching consequences for our understanding of the regulation of rhizosphere processes. While the assembly of rhizosphere bacterial communities is bottom-up controlled through the resource supply from root metabolites, our results support the hypothesis that the net effect may depend on the strength of top-down control by protist grazers, thereby stabilizing the functional performance of bacterial microbiomes on plant surfaces.

## Acknowledgements

This work was supported by the DFG projects (SM59/11-1/GR568121) and the Priority Program “Rhizosphere Spatiotemporal Organisation” (SPP 2089) of the German Science Foundation (DFG), as well as the Cluster of Excellence on Plant Sciences CEPLAS (EXC 1028). The authors declare no conflict of interest.

